# Temporal modulation of host aerobic glycolysis determines the outcome of *M. marinum* infection

**DOI:** 10.1101/507160

**Authors:** Meng Lu, Lingling Xie, Yuanqing Kan, Lixia Liu, Dong Wenyue, Jintao Feng, Yuchen Yan, Gang Peng, Mingfang Lu, Chen Yang, Chen Niu

## Abstract

Macrophages are the first-line host defense where the invading *Mycobacterium tuberculosis* (Mtb) encounters. It has been recently reported that host aerobic glycolysis was elevated post the infection by a couple of virulent mycobacterial species. However, whether this metabolic transition is required for host defense against intracellular pathogens and the underlying mechanisms remain to be further investigated. By analyzing carbon metabolism, we found that macrophages infected by *M. marinum*, a surrogate mycobacterial specie to Mtb, showed a strong elevation of glycolysis. Next, three glycolysis inhibitors were examined for their ability to inhibit mycobacterial proliferation inside RAW264.7, a murine macrophage-like cell line. Among them, a glucose analog, 2-deoxyglucose (2-DG) displayed a protective effect on assisting host to resist mycobacterial infection, which was further validated in zebrafish-infection model. The phagocytosis of *M. marinum* was significantly decreased in macrophages pre-treated with 2-DG at concentrations of 0.5 and 1 mM, at which no inhibitory effect was posed on *M. marinum* growth *in vitro.* Moreover, 2-DG pre-treatment exerted a significant protective effect on zebrafish larvae to limit the proliferation of *M. marinum,* and such effect was correlated to tumor necrosis factor alpha (TNF-α). On the contrary, the 2-DG treatment post infection did not restrain proliferation of *M. marinum* in WT zebrafish, and even accelerated bacterial replication in TNF-α^-/-^ zebrafish. Together, modulation of glycolysis prior to infection boosts host immunity against *M. marinum* infection, indicating a potential intervention strategy to control mycobacterial infection.

**Author Summary:** As an intracellular pathogen, Mtb exploits multiple strategies to invade and hijack macrophages for its own advantages. Accordingly, recent investigations have shown that Mtb infection is accompanied with an alteration of host glucose metabolism. Macrophage and zebrafish infection models of *M. marinum*, facilitating our understanding towards mycobacterial pathogenesis, were applied in this study. We found that the pre-treatment of macrophages with a glucose analog, 2-DG, inhibited aerobic glycolysis and made host cells more inert to phagocytose the bud. In infected zebrafish larvae, bacterial load inside host pretreated with 2-DG remains at a significantly lower level compared to the untreated group. These findings imply that the modulation of host glycolysis regulates the fate of *M. marinum* infection, and indicate a promising metabolic target in TB intervention.

## Introduction

Tuberculosis (TB), a serious chronic infectious disease caused by *Mycobacterium tuberculosis* (Mtb), is still a major threaten to public health worldwide. The interaction between Mtb and macrophages was initiated with phagocytosis [1]. Downstream cell defense events will then be switched on, including phagosome maturation, acidification, the fusion between phagosome and lysosome. Besides, cytokines including tumor necrosis factor alpha (TNF-α) and interleukin 1 beta (IL-1β) secreted by infected macrophage cells play important roles mediating host to fight against Mtb infection. Meanwhile, autophagy plays an interesting role in this interaction [2,3]. However, it has been illustrated from numerous studies that, through the expression of various virulent factors, Mtb could dampen these host immune responses and survive the battle [4,5]. The altered immunometabolism of macrophage was observed post mycobacterial infection [6]. Among these alterations, the elevation of glycolytic products has been noticed from a few recent studies [7,8]. Meanwhile, glycolytic pathways were dramatically induced at transcriptional level [9]. However, the influence of altered host glycolysis in TB pathogenesis remains to be understood. In this study, *M. marinum* [10], a surrogate mycobacterial specie to Mtb was utilized. We firstly investigated the carbon metabolism shift and immune responses of macrophages post *M. marinum* infection. The glucose uptake and lactate secretion of murine macrophage-like cell line RAW264.7 (RAW cells) were significantly boosted post infection, and glycolysis was elevated in infected macrophages. Next, three glycolysis inhibitors were tested for their effects on the replication of intracellular bacteria, and only 2-DG exerted protective effects via modulating macrophage phagocytosis and following immune responses.

Furthermore, the protective effects of 2-DG were validated *in vivo* using *M. marinum*-zebrafish infection model, which is widely used to study the role of host immunity in mycobacterial pathogenesis [11,12,13]. Specifically, previous studies have revealed the crucial role of aerobic glycolysis in cytokines production [14,15,16]. We herein found that TNF-α, an important pro-inflammatory cytokine in TB immunity [17,18,19], was boosted in mouse peritoneal macrophages upon the 2-DG pre-treatment. Moreover, 2-DG was applied in both WT and TNF-α^-/-^ zebrafish [20] prior to or upon *M. marinum* infection, and bacterial proliferation was measured at various time points post infection. This study herein revealed that serial immune responses were significantly enhanced by 2-DG pretreatment. In addition, the understanding on the role of host glycolysis in mycobacterial infection may provide a novel angle on identifying host metabolic targets in TB intervention.

## Results

### Macrophages displayed an elevated glycolysis during *M. marinum* infection

To elucidate the carbon metabolism shift of macrophages upon and post *M. marinum* infection (Fig 1A), glucose uptake and lactate secretion of macrophages post *M. marinum* infection was firstly determined. A decline of glucose concentration in culture medium was observed on 24 hour post infection (hpi), which was even more dramatic on 48 hpi, indicating a significant glucose uptake of RAW cells following *M. marinum* infection (Fig 1B). Meanwhile, the concentration of lactate in cell culture medium remarkably increased in infected macrophages compared to the uninfected control on both 24 hpi and 48 hpi. In order to prove that the utilization of glucose is via glycolysis, RAW cells were lysed at the same time points, and the glycolytic intermediates were measured. Among them, an increase of fractional labeling for glycolytic intermediates including hexose-6-phosphate, 3-phosphoglyceric acid, and pyruvate (Fig 1C) was observed. Together, glucose uptake, lactate secretion, and the glycolysis were remarkably elevated in RAW cells post *M. marinum* infection. To determine if increased glycolysis affects the downstream TCA cycle, the relative concentration of TCA intermediates were calculated. Compared to the uninfected group, an increased concentration of TCA intermediates (S1 Fig) was observed, and we speculated that the TCA flux might increase following the elevated glycolysis.

**Fig 1.**
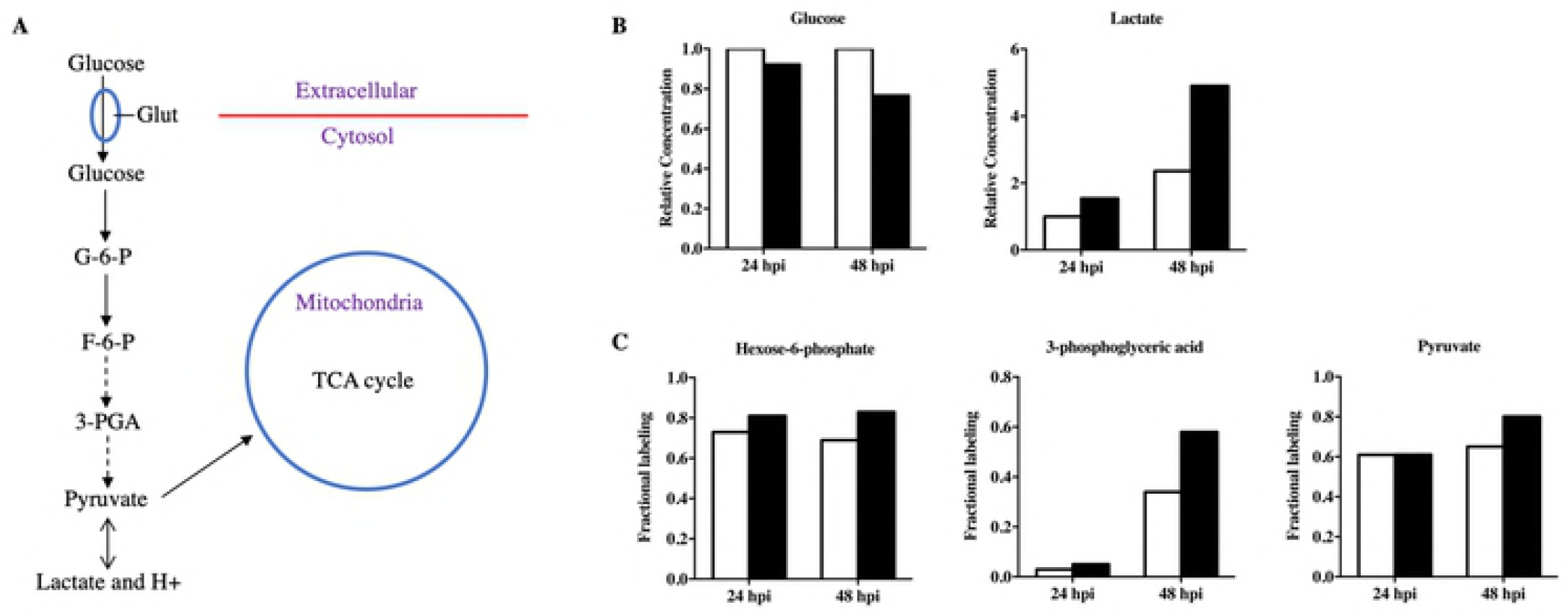
The increased glucose uptake, lactate secretion and glycolysis of RAW264.7 cells post *M. marinum* infection. (A) Sketch of glucose uptake and glycolysis pathway. (B) The relative concentrations of glucose and lactate in cell culture were measured by HPLC at 24 hpi and 48 hpi (black bar), uninfected control is depicted as white bar. The degree of glucose uptake was calculated as the reduction of glucose concentration in cell culture medium, which was normalized to the original concentration (4.5 g/L) in C-DMEM. (C) The fractional labeling (FL) of several other glycolytic intermediates were calculated in lysed RAW264.7 cells cultivated in KO-DMEM plus [U-13C] glucose as the sole glucose source.

### Glycolysis inhibitor 2-DG reduced *M. marinum* burden inside macrophages through phagocytosis inhibition

To further address the consequence of enhanced glycolysis of RAW cells upon *M. marinum* infection, several glycolysis inhibitors were added separatively or in combination into cell culture medium at the time of infection. The inhibitors selected were oxamate, 2-deoxy-D-glucose (2-DG), and sodium dicholroacetate (S2A Fig), which target several key enzymatic steps of glycolysis including lactate dehydrogenase, hexokinase, and pyruvate dehydrogenase kinase separately. Among them, only 2-DG displayed a remarkable effect inhibiting the bacterial burden inside macrophages (S2B Fig). In addition, the *in vitro* growth of *M. marinum* in 7H9O broth medium was not inhibited by 2-DG at the concentrations less or equal to 1mM (Fig 2A), but the growth of *M. marinum* was suppressed when 2-DG concentration was equal or higher than 5 mM.

**Fig 2.**
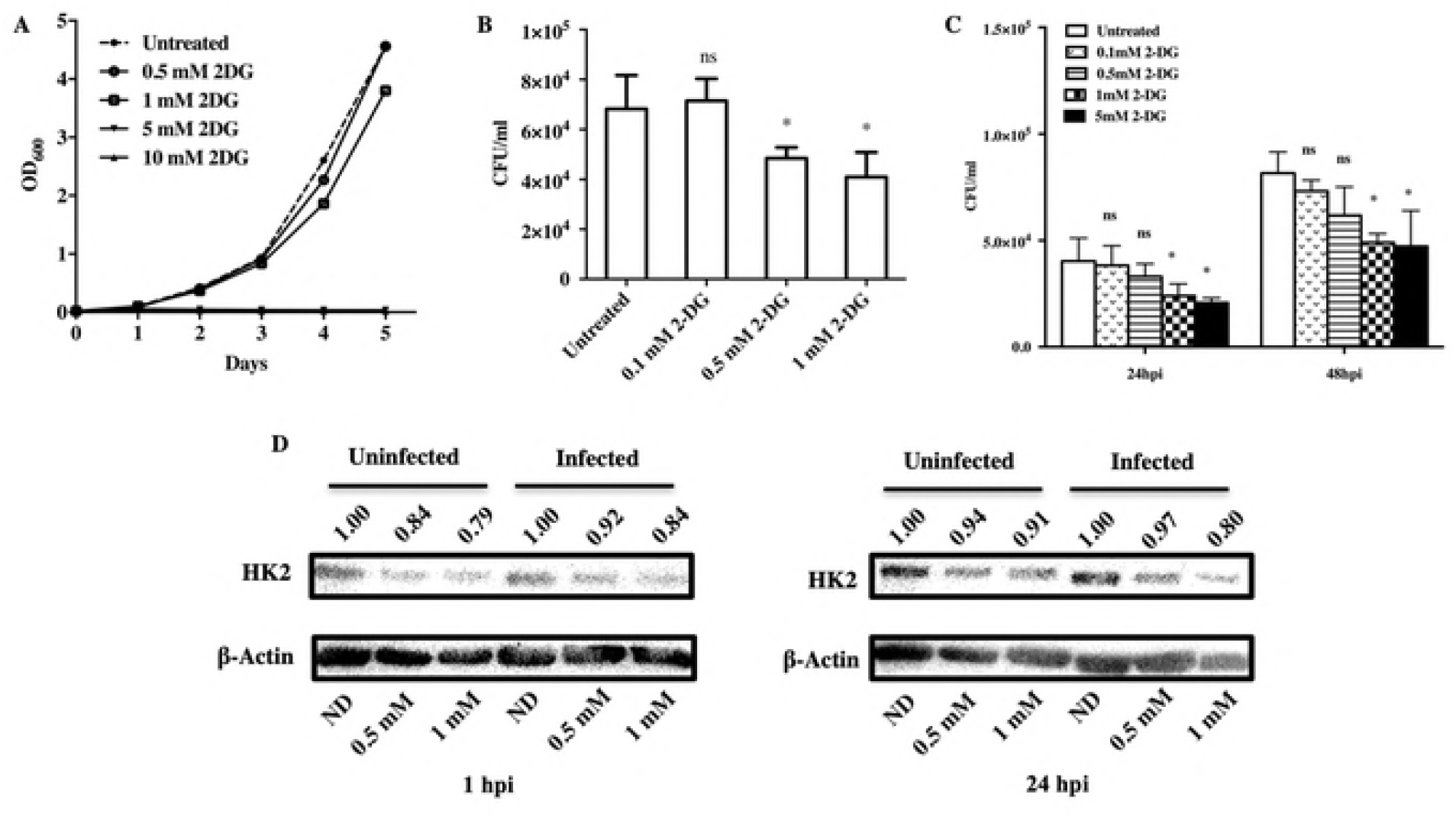
2-DG pretreatment inhibited phagocytosis of *M. marinum* by RAW264.7 cells. (A) The *in vitro* growth of *M. marinum* in 7H9+OADC broth containing various concentrations of 2-DG. (B-C) 1 × 10^5^ RAW264.7 cells untreated or pretreated with 2- DG (0.1, 0.5, 1 and 5mM) were infected by *M. marinum* at MOI = 10. At various time points post infection, RAW264.7 cells were rinsed and lysed, intracellular bacterial load of *M. marinum* load was measured by spreading onto 7H10+OADC agar plates. For statically analysis, Student’s T test was performed between 2-DG pretreated groups and the untreated group. *, p<0.05, “ns”, no significant difference. (B) 24 hpi, (C) 48 hpi. (D) Relative HK2 protein levels in RAW264.7 cells were determined by western blot analysis.

The decreased bacterial load of *M. marinum* recovered in 2-DG treated macrophages might be caused by either the reduced phagocytosis or enhanced bactericidal ability of macrophages. First, the effects of 2-DG pretreatment on phagocytosis of RAW cells were examined, and the inhibitory effects of 2-DG on phagocytosis were dose-dependent (Fig 2B). When RAW cells were cultivated in medium containing 0.5 mM and 1 mM concentrations of 2-DG, 72.9± 15.1 % and 624 ± 20.0 % bacilli were phagocytized relative to the untreated group. The tendency of such inhibition was maintained at later time points (24 and 48 hpi) as shown in Fig 2C. Hexokinase 2 (HK2) is the first enzyme in glycolysis which phosphorylates glucose to produce glucose-6-phosphate (G6P). The glucose analogue 2-DG is phosphorylated by HK2 to 2-DG-P but cannot be further metabolized, thus causes the metabolic block which inhibits glycolysis [21]. Accordingly, we found that the expression of HK2 was slightly increased in RAW cells post *M. marinum* infection, which was inhibited by 2- DG pre-treatment (Fig 2D).

### 2-DG pretreatment led to autophagy or apoptosis of RAW cells in a dosage-dependent manner

To determine cellular responses that 2-DG pretreatment might pose on, autophagy and apoptosis, two pathways among major strategies by which macrophages combat with *M. marinum* were investigated. Autophagy begins with the formation of double-membrane autophagosome, following with autophagolysosome development which could be marked by Cyto-ID. Raw cells were treated 24h and 48h with 2-DG at concentrations of 0, 1, 5 or 10 mM, followed by FACS analysis (Fig 3A). The treatment of RAW cells with 1 mM 2-DG for 24h and 48h strongly induced cellular autophagy (Fig 3B), which was further aggravated when higher concentration of 2-DG was applied. During autophagosome formation, the cytosolic form of the microtubule-associated protein 1A/1B-light chain 3 (LC3-I) is converted to lipid-bound LC3-II. Consistently, positive LC3-II punctative pattern was observed in 2-DG pretreated RAW cells under confocal microscope (Fig 3C and 3D). Furthermore, increased apoptosis was observed only when RAW cells received 10 mM 2-DG on both 24h and 48h, indicating that cell apoptosis was dependent on a relatively high concentration of 2-DG (S3 Fig). Together, these data indicates that autophagy was induced by a 24h or prolonged treatment of 2- DG, which might contributes to earlier reduction of intracellular bacilli post *M. marinum* infection.

**Fig 3.**
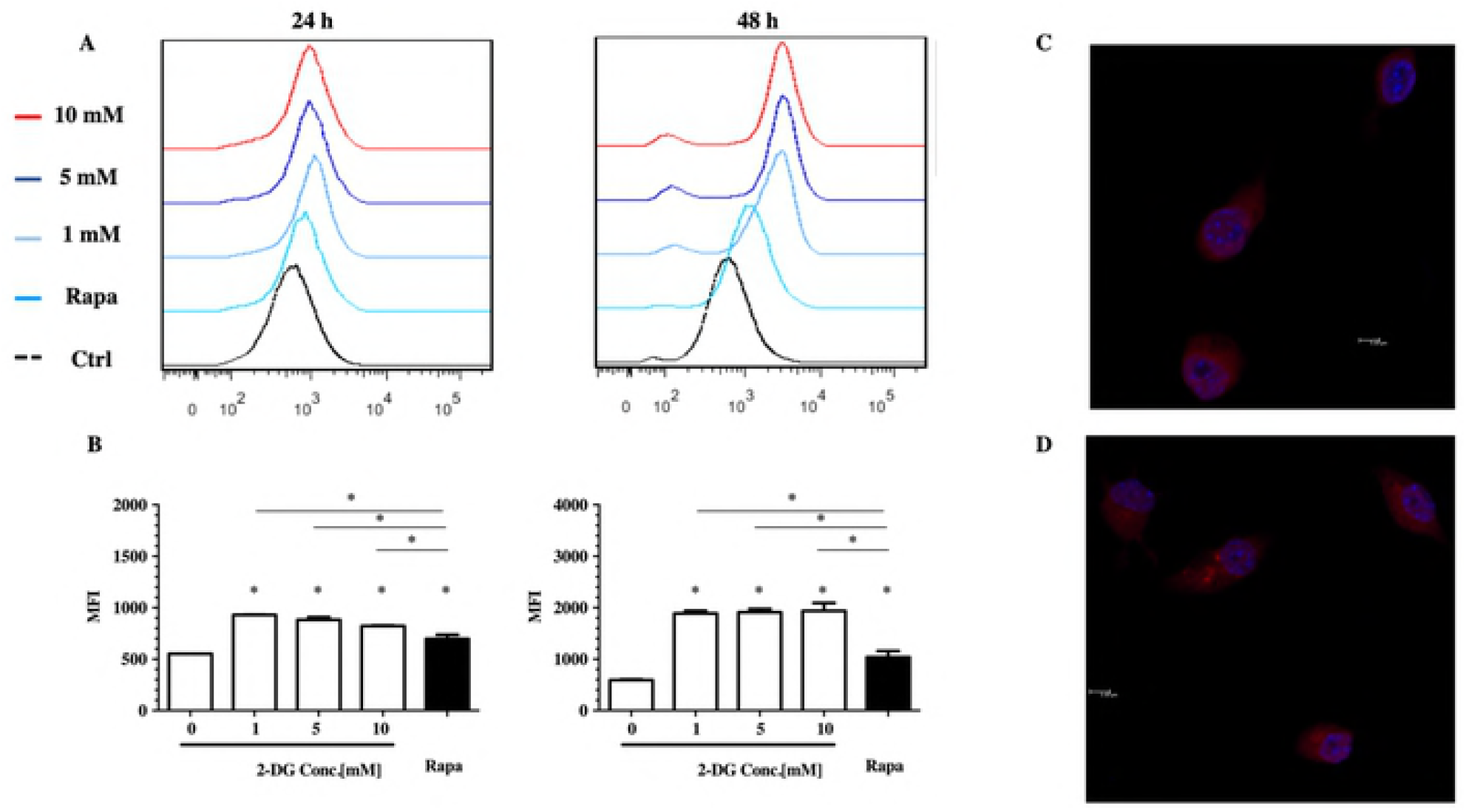
Effect of 2-DG treatment on autophagic death. (A) Representative flow cytometric dot plots illustrating autophagy of Raw264.7 cells treated with 2-DG at 0, 1, 5 and 10 mM for 24h and 48h. 1 μM Rapamycin applied in Cyto-ID^®^ Kit was used as positive control. (B) Bar graphs presenting the degree of autophagy treated with 2-DG at 0, 1, 5 and 10 μM for 24 h and 48h. Cells were treated with green fluorescent Cyto-ID^®^ to detect autophagic vacuoles and subjected to flow cytometric analysis. Data are the mean ± SD values of three independent experiments. **P<0.05 by two-tailed t test. (C&D) 2-DG pretreatment induced autophagy in RAW264.7 cells. Cell nuclei were stained using Hochest dye and the red fluorescence marked LC3-II which aggregates during autophagy in cytoplasm. (C) Control group (without 2-DG) and (D) Cells treated in DMEM with 1mM 2-DG for 24 hours.

### 2-DG pre-treatment inhibited the proliferation of *M. marinum* in WT zebrafish

To examine if 2-DG also exerts protective effects *in vivo,* the *M. marinum-* zebrafish infection model was utilized. A volume of 1 nL 2-DG at concentrations of 1 or 5 mM was injected into one-cell stage zebrafish embryo on 0 hpf (hour post fertilization). On 28 hpf, *M. marinum* carrying a plasmid pTEC15 expressing green fluorescent protein (Mm pTEC15) was injected into the caudal vein at a dosage of 1000 CFU per fish (Fig 4A). To evaluate the diffusion capability of 2-DG, a fluorescent analog 2-NBDG, was injected and an even diffusion has been observed at 28 hour post fertilization (hpf) when the infection initiated. Meanwhile, no growth defect of zebrafish larvae was observed in 2-DG injected WT zebrafish larvae (data not shown). The pre-treatment of 2-DG exerted a significant protective effect for host to resist *M. marinum* proliferation lasting till 5 dpi, the end time point of experiments (Fig 4B-4D). We next asked if such protective effects of 2-DG could be achieved through other treatment methods. To our surprise, protective effects were not observed either when 2-DG was injected with Mm pTEC15 simultaneously (Fig 5C) or immersed into embryo medium (S4 Fig), whereas rifampicin displaying a known effect to inhibit the proliferation of *M. marinum* in zebrafish as reported [22].

**Fig 4.**
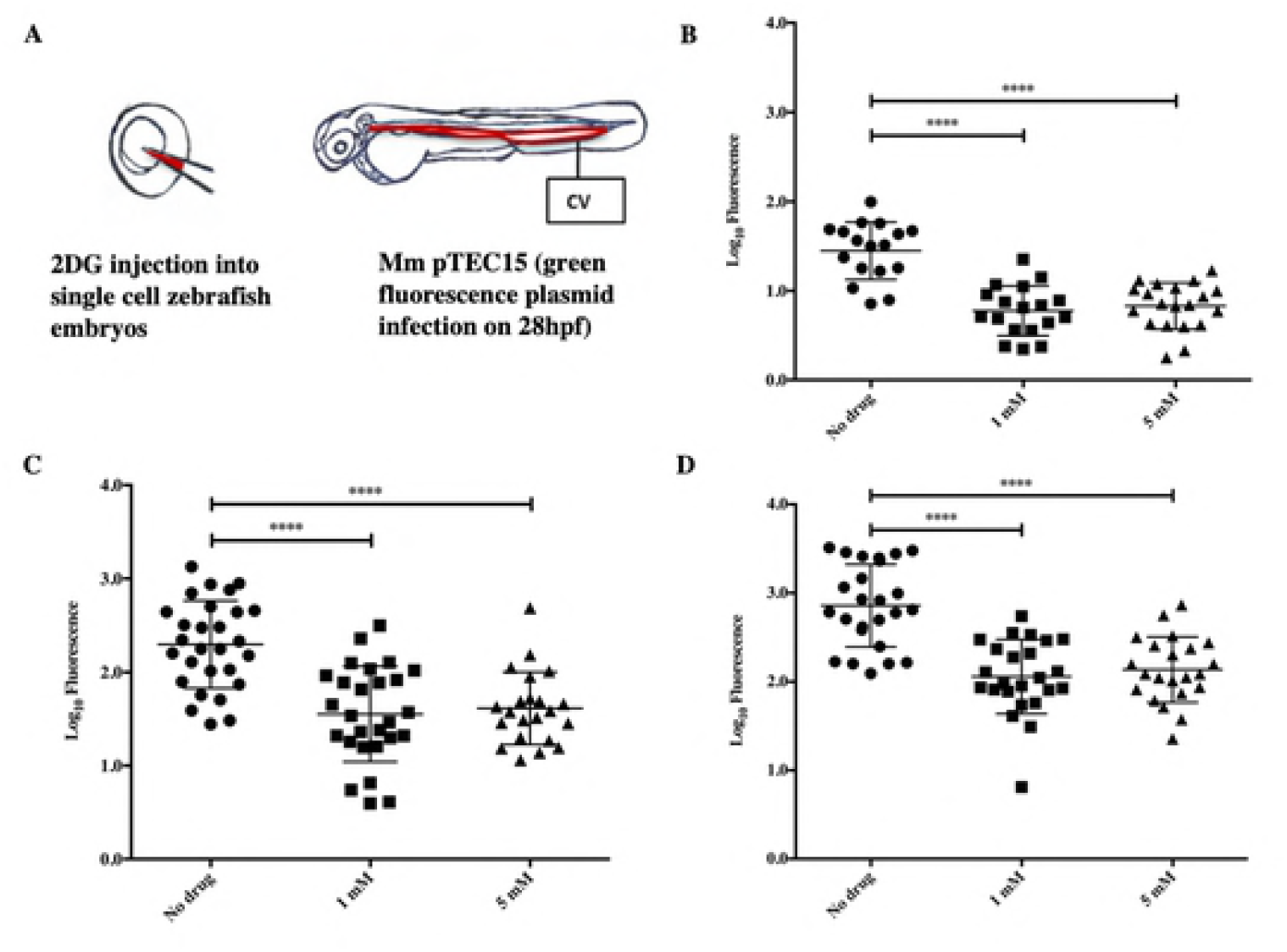
*M. marinum* proliferation *in vivo* was inhibited by 2-DG pretreatment. An initial infection dosage was 1000 colony forming unit (CFU) Mm pTEC15 per zebrafish larvae. (A) The diagram of 2-DG injection on 0 hpf and infection of larvae with Mm pTEC15 on 28 hpf. (B-D) Bacterial load in zebrafish larvae were measured on various time points post infection. Zebrafish were imaged using AMG EVOS fluorescence microscope, and bacterial load were analyzed by counting fluorescence pixels of images using software Image J. (B) 1 dpi, (C) 3dpi, and (D) 5 dpi.

### 2-DG accelerated *M. marinum* proliferation in zebrafish missing TNF-α

To understand how 2-DG pretreatment facilitates host to inhibit *M. marinum* proliferation, the transcriptional level of several crucial cytokines related to TB immunity including TNF-α, IL-6, IL-10 were examined in zebrafish larvae (data not shown). Among them, the transcriptional level of TNF-α was significantly elevated in 2-DG pretreated zebrafish larvae on 28 hpf (Fig 5A). Consistently, 2-DG pretreatment had also boosted the secretion of TNF-α in isolated mouse peritoneal macrophages stimulated with LPS, compared to PBS as a mock (Fig 5B).

**Fig 5.**
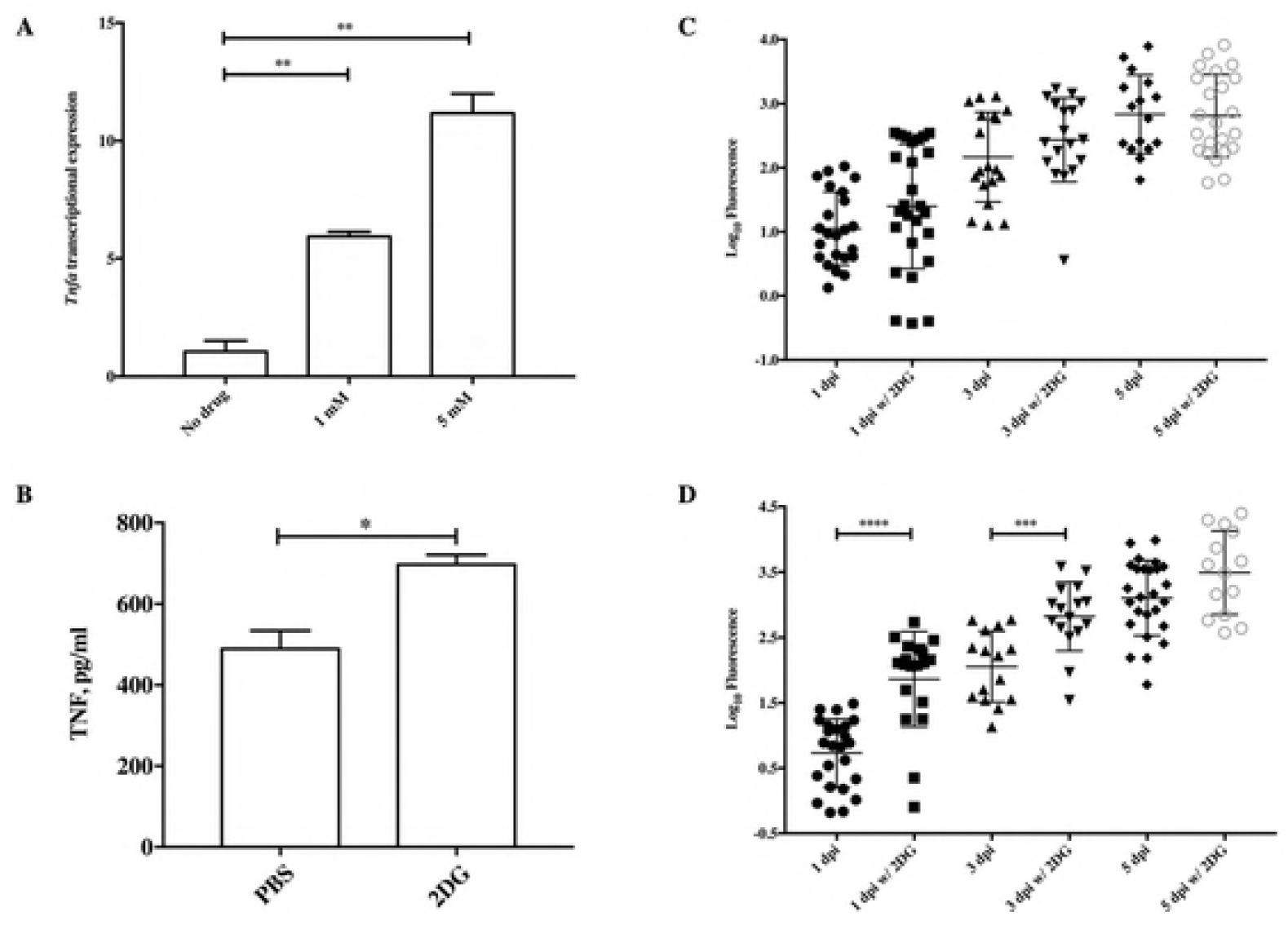
2-DG accelerates *in vivo* proliferation of *M. marinum* in zebrafish lack of TNF-α. (A) The transcription of TNF-α in zebrafish larvae on 28 hpf 2-DG pretreatment. (B) The 2-DG pretreatment augments TNF-α secretion by mouse peritoneal macrophages cultured in medium containing LPS. (C) WT zebrafish or (D) TNF-α^-/-^ zebrafish larvae was infected on 28 hpf by CV injection with a mixture of 2-DG (5 mM) and 200 CFU Mm pTEC15, and bacterial load in zebrafish larvae were measured by counting fluorescence pixels of images using software Image J.

To demonstrate whether 2-DG protective function in *vivo* is TNF-α dependent, TNF-α^-/-^ fish previously constructed [20] and maintained in our laboratory was utilized. Due to the high mortality ratio of TNF-α^-/-^ larvae post 2-DG injection on single cell stage after multiple trials (data not shown), 2-DG pretreatment in TNF-α^-/-^ larvae could not be performed. Thus, *M. marinum* and 2-DG were co-injected into TNF-α^-/-^ larvae on 28 hpf, with WT zebrafish larvae as a control. Intriguingly, 2-DG did not exhibit protective effects in WT larvae under such experimental condition (Fig 5C), and accelerated *M. marinum* proliferation in TNF-α^-/-^ larvae instead (Fig 5D). Thus, 2-DG could not exert protective effects once *M. marinum* had already infected zebrafish larvae. It suggests that 2-DG posed detrimental effects when TNF-α is missing in host.

## Discussion

Emerging evidence have indicated that glucose metabolism of host cells altered post infections by several virulent mycobacterial species. For example, glucose uptake with a concomitant increase in glucose-6-phosphate dehydrogenase activity was observed in Schwann cell infected by *M. leprae* [23]. Along with this, the surface localization of glucose transporters GLUT1 and GLUT3 increased [24]. In addition to the elevated glucose uptake, macrophage glycolysis was elevated post infection by either Mtb [8] or *M. avium* [25]. In present study, both the augmented glucose uptake and elevated glycolysis in RAW cells infected with *M. marinum* were observed, accompanied with a significant increase of lactate secretion (Fig 1). It indicates that *M. marinum* may apply similar strategies to modulate glycolysis of macrophages.

The present study further characterized the consequence of glycolytic modulation in *M. marinum* infection using 2-DG, both *in vitro* and *in vivo.* In RAW cells, 2-DG made macrophages more inert to mycobacterial invasion (Fig 2B). A few decades ago, it had been noticed that 2-DG selectively inhibits Fc and complement receptor-mediated phagocytosis in mouse peritoneal macrophages [26]. In addition, our findings that autophagy was induced by the pre-treatment with 1 mM 2-DG (Fig 3) indicated that it may assist host cells against mycobacterial infections in multiple means. Phagocytosis and autophagy are both ancient, highly conserved processes respectively involved in the removal and destruction of organisms in the cytosol. In support of our findings, effects of 2-DG on several other intracellular bacteria has been studied over the past two decades.

An early study indicates that 2-DG induced metabolic stress on mice to resist the infection with *Listeria monocytogenes* [27]. Later on, it was found that 2-DG facilitates the autophagy of A/J mouse macrophages, and suppressed the intracellular multiplication of *Legionella pneumophila* [28]. In line with it, HK2 integrates glycolysis and autophagy to confer cellular protection [29], and enzymatic disruption might be a novel strategy for treating infecting pathogens [30]. Intriguingly, high concentration of 2-DG (>5 mM) was found to induce apoptotic death of RAW cells (S3 Fig). Since autophagy can be pharmacologically modulated, it is considered as one of therapeutic opportunities for TB [3]. The potential effects of 2-DG and role of HK2 in TB pathogenesis are worthy to be further explored.

Previous studies using zebrafish model confirmed that TNF-α is indispensable for controlling the proliferation and dissemination of *M. marinum* [17]. It was illustrated from this study that protective effects of 2-DG is correlated with TNF-α (Fig 5C and 5D). Hence, we speculated that there is cross-talk between 2-DG and TNF-α mediated cell pathways. It has been demonstrated that the elevated aerobic glycolysis is beneficial for macrophage cells and/or host to combat with intracellular Mtb [7], and Mtb could dampen the levels of proinflammatory IL-1β and increased anti-inflammatory IL-10 via influencing host cell glycolysis. Additional evidence came from a recent study that glycolysis inhibition using 2-DG posed detrimental effects on host immunity post *Mtb* infection of mice [31]. In line with this, 2-DG injection along with *M. marinum* infection of WT zebrafish did not inhibit bacterial proliferation *in vivo* (Fig 5C). However, given other findings in this study that 2-DG pre-treatment improve both macrophages and zebrafish to resist the invasion and proliferation of *M. marinum* (Figs 2-4), we speculate that glycolysis might play double-edge sword effects during mycobacterial pathogenesis. At the initial stage of infection, virulent mycobacteria might hijack the glycolysis for its invasion into macrophages. Later on, elevated glycolysis is correlated with boosted host immunity to fight against the bug. It remains to be elucidated that to which degree these conflicting effects mediate the interaction between the bacilli and macrophages. A well-known virulence factor, Esat6 was found to induce metabolic flux perturbations to drive foamy macrophage differentiation [32]. Together with the observation that *Mtb* could utilize various carbon sources inside host [33], it implies that the altered host glycolytic pathways might be the outcome of the complicated interaction between pathogenic mycobacterium and host. The present study implies that timing may be a key factor to determine whether *M. marinum* or macrophages benefit from the elevated glycolysis during infection.

With the increase of multi-drug resistant (MDR) strains of Mtb and slow progress of developing new antimicrobials for treating TB, an emerging concept for treating TB is host-directed therapy (HDT) [34,35,36]. For example, metformin, a drug used in treating diabetes, has been validated as a candidate with high potential against TB [37]. Key enzymes involved in glycolysis might be attractive targets for anti-TB drug screening, in which case glycolysis inhibitors might be potential candidates in HDT, since 2-DG has already been tested for safety in multiple clinical trials [38]. Our study underscores the importance of glycolysis in TB pathogenesis, and further study on this complex interaction is a prerequisite for developing novel HDT strategies for TB.

## Material and methods

### Strains and culture conditions

*M. marinum* M strain (ATCC BAA-535) was routinely cultivated in Middlebrook 7H9 broth or on 7H10 agar enriched with 10% OADC (oleic acid-albumin-dextrose-catalase) and 0.4% volume/volume (v/v) glycerol. When necessary, 50 μg/ml of hygromycin was included to maintain *M. marinum* carrying pTEC15, a plasmid carrying GFP under a mycobacterial promoter [22]. For growth measurement, strains were cultured in 7H9 broth with or without 2-DG. In addition, 0.02% v/v tyloxapol was added into 7H9 broth to reduce bacterial clumping.

### Macrophage culture conditions and compounds

Dulbecco’s Modified Eagle Medium containing 4.5 g/L glucose (Gibco) and 10% FBS (C-DMEM) were used to cultivate RAW cells. Cells were infected by using single cell *M. marinum* at a multiplicity of infection (MOI) =1 for 5 hours at 32 °C. Then, extracellular bacteria were killed using gentamycin at a concentration of 200 μg/ml for 2h, then fresh DMEM were replaced. For phagocytosis assay, RAW cells were infected by using single cell *M. marinum* at an MOI=10 for 4 hours at 32 °C. Chemicals 2- Deoxy-D-glucose, oxmate, and DCA were purchased from Sigma.

### Metabolite measurements

The absolute concentration of metabolites was measured on 24 hours post infection (hpi) and 48 hpi. Glucose and lactate in cell culture medium were quantitated by high pressure liquid chromatography (HPLC) using an Agilent model 1260 instrument equipped with a Shodex RSpak KC-811 Column (8 × 300 mm; Shodex) and a UV detector (Agilent) operated at 210 nm. The mobile phase solution was 6 mM HClO_4_ in water and pumped at a flow rate of 1.0 ml/min, the temperature of the column was kept at 50 °C.

For the measurement of celluar metabolites, RAW cells were centrifugated and cell pellets were resuspended in 2 ml of 80:20 (v/v) methanol/water precooled to −80 °C separately and lysed by mechanical vortexing. After a centrifugation for 3 min at 16200 ×g, the supernatants were collected and analyzed by UPLC system (Waters) coupled to a Q Exactive hybrid quadrupole–orbitrap mass spectrometer (Thermo Fisher). Injection volume was 10 μl. Solvent A was 50 mM ammonium acetate adjusted to pH 9.0 with ammonium hydroxide, and solvent B was acetonitrile. Metabolites were separated with a Luna NH2 column (100 mm × 2 mm, 3-μm particle size; Phenomenex). The column was maintained at 15°C with a solvent flow rate of 0.3 ml min-1, and the gradient of B was as follows: 0 min, 85%; 3 min, 30%; 12 min, 2%; 15 min, 2%; 16 min, 85%; 23 min, 85%. The mass spectrometer with a heated electrospray ionization source was operated in negative modes, and the key parameters were same as described in [39]. Data were analyzed using the Xcalibur software.

For [U-13C] glucose labelling experiments, the fractional labeling (FL) of different metabolites was calculated according to a published protocol [40] as following:

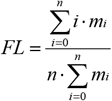

where *n* represents the number of C atoms in the considered fragment and *i* the different mass isotopomers, *m_i_* represents the amount of the compound with i×^13^C atom.

### Zebrafish Infection

AB wild type (WT) and TNF-α^-/-^ mutant zebrafish [17] were utilized in this study. For larval infection, experiments were performed as described in a recent paper [41]. Regarding the dosage of 2DG injection, 1 nL of 2-DG at 10 mM concentration at the single cell level of zebrafish larvae is approximately 50 mg/kg body weight, which is in the safety range among multiple clinical trials [38].

### Animal Ethics Statement

All experiments using zebrafish in this study was adhered to the protocol, which was reviewed and approved by Institutional Animal Care and Use Committee (IACUC) of Fudan University. The approval and identification number is # 20120105-001, and the protocol is adhered to the regulations/guidelines by Office of Laboratory Animal Welfare, NIH.

Briefly, zebrafish are euthanized in a manner that minimizes their discomfort, pain, and the time to death. Fish are euthanized by an excess dose of buffered MS-222 (tricaine) on ice, 150-250 ppm (milligrams per liter), depending on the animals’ size, age and duration of anesthesia. The zebrafish are placed in an immersion both with tricaine, and placed on ice. If no movements are observed after 15 minutes, the euthanasia is complete. This method is consistent with the recommendations of the AVMA Guidelines on Euthanasia. The dead or euthanized fish and their waste water will be disinfected with a 10% bleach solution (final concentration) or a 2% alkaline glutaraldehyde solution for a minimum of 30 minutes. The dead or euthanized fish are placed in specific labeled biohazards bags and disposed in biohazards waste containers for pick-up by the biohazards waste disposal company that is contracted by Fudan University.

### Imaging of macrophages and zebrafish

Zebrafish were imaged under FITC channel of AMG EVOS fluorescence microscope. Bacterial burdens of *M. marinum* were determined in larvae by measuring fluorescence pixel counts (FPC) through the ImageJ software. For immunostaining of autophagy marker LC3-II in RAW cells, fluorescence signals were detected by using confocal laser scanning microscopy (CLSM) Leica SP8 at an excitation wavelength of 375 and 543 nm separatively. Images were processed using LASAF Lite software.

### FACS Assay

Raw cells were cultured in C-DMEM contain different concentration of 2-DG (0, 1, 5 and 10mM). For apoptosis analysis, at 24h or 48h, cells of each treatment group were collected by repeated blow and wash once by C-DMEM. After a quick spin (500g, 5min), Raw cells were resuspended in 100ul Binding buffer, then 50ul antibody solution contain 0.5ul Annexin V FITC (1/100) and 1ul 7AAD PerCP-Cy5.5 (1/50) was added followed by incubating for 20min in dark. Finally, wash cells for two times using MACS and resuspend cells in 200ul FACS to perform the FACS assay. Cell apoptosis was calculated by the percentage of Annexin V+ 7AAD-.

To determine the autophagy of Raw cells after 2-DG treatment, fluorescent Cyto-ID^®^ that can stain autophagic vacuoles was obtained from Enzo Life Sciences Inc. (Farmingdale, NY, USA). Autophagy detection was performed according to the product manuscript. In brief, at appropriate time point, each sample was washed by DPBS, then cultured in 200ul C-DMEM containing 0.2ul Cyto-ID Green containing indicator and incubate at 37C, 5% CO2 in the dark. The Cyto-ID containing medium was wash away with DPBS. Using cold DPBS containing 2% FBS, cells were resuspend and cell autophagy analysis was performed through FACS. The mean fluorescence intensity (MFI) was calculated.

## Acknowledgments

The plasmid pTEC15 was a gift from Lalita Ramakrishnan, University of Cambridge, UK. We are grateful to insightful advice from Dr. Liangdong Lyu at Fudan University.

## Supporting information

**S1 Fig. Elevated TCA cycle of macrophages post *M. marinum* infection.**

The relative abundance of each TCA cycle intermediate was calculated by summing of peak areas of all isotopomers acquired by LC-MS, including (A) citrate, (B) aconitate, (C) itaconate, (D) isocitrate, (E) α-KG (α-ketoglutarate), (F) succinate, (G) fumarate, and (H) malate. Infected macrophages (Black Bar), the uninfected group (White Bar).

**S2 Fig. Intracellular replication of *M. marinum* is significantly inhibited by 2-deo-D-glucose.**

(A) Metabolic targets of selected inhibitors. (B) Intracellular bacterial load of *M. marinum* was calculated on 0, 24, 48 and 72hpi in infected macrophages. Untreated (U) or treated with Oxamate (Ox), 2-deo-D-glucose (2-DG) or sodium dicholroacetate (SD) alone and combination with various drugs.

**S3 Fig. 2-DG induced apoptotic death in Raw264.7 cells in a concentration - dependent manner.**

(A) Representative flow cytometric dot plots showing the percentage of specific cell populations (live, early apoptosis, and late apoptosis) in Raw264.7 cells treated with 2DG at 0, 1, 5 and 10 mM for 24 h and 48h. (B) bar graphs showing the percentage of apoptotic cells in Raw264.7 cells treated with 2DG at 0, 1, 5 and 10 mM for 24 h and 48h. Cells were double stained using annexin V:FITC and 7-AAD PerCP-Cy5.5 to detect cells undergoing early apoptosis (left) and late apoptosis (right). Data are the mean ± SD of three independent experiments. **P<0.05 by two-tailed t test.

**S4 Fig. The *in vivo* proliferation of *M. marinum* was not inhibited by 2-DG added into embryo medium.**

WT zebrafish was infected on 28 hpf by caudal vein injection with an Initial dosage of 200 cfu per fish, and bacterial load of Mm pTEC15 in zebrafish larvae were measured by counting fluorescence pixels of images using software Image J. (A) 1 dpi, (B) 2 dpi, (C) 3 dpi, (D) 4 dpi.

